# A dynamical systems model of arousal-driven behavioural state transitions

**DOI:** 10.1101/2025.10.31.685593

**Authors:** Philippa A. Johnson, Sander Nieuwenhuis, Jorge Mejías, Anne E. Urai

## Abstract

When completing a task, animals switch between being engaged and disengaged. What neural and physiological processes trigger these behavioural state transitions? We estimated the engagement state of mice performing a visual decision-making task using a hidden Markov model of response times and found that intermediate arousal, as measured using baseline pupil size, was associated with more engagement in the task. Additionally, we show that changes in arousal (both mean and variability of the pupil baseline) predict changes in behavioural state. To explain this, we propose a double-well model, in which arousal causes behavioural state transitions by reshaping the attractor landscape of population neural activity. These results highlight a possible mechanism of arousal-related changes in behaviour.

## Introduction

Perceptual decision-making, the process of transforming sensory input into choices, is an inherently variable process: behaviour differs when making repeated decisions about the same sensory evidence (Duffy et al., 2025; Urai, 2025). One source of variability in decision-making is changes in behavioural state. During visual decision-making, animals frequently disengage during sessions of decision-making, showing periods of inaccurate and biased responses (Ashwood et al., 2022; Hulsey et al., 2024). There is evidence that humans spend thirty to fifty percent of their waking hours distracted or mind wandering (Killingsworth & Gilbert, 2010), similarly corresponding to periods of low accuracy and slow responses (Zhang & Kool, 2025).

Neural correlates of being engaged or disengaged in a task have been well-characterised (Jacobs et al., 2020; Mittner et al., 2016; Rothlein et al., 2018; Steinmetz et al., 2019), so, here, we focus on the question of what neurophysiological mechanisms drive the transitions between these states. Subcortical, neuromodulatory arousal systems may play a key role. Neuromodulatory activity can cause a variety of behavioural state transitions, including sleep-wake (Silverman et al., 2025), explore-exploit (Kane et al., 2017), and different task-related states within a trial (Corbetta et al., 2008; Jahn et al., 2020; Sara & Bouret, 2012). A recent study identified a ‘subcortical switchboard’, whereby activity of serotonin neurons in the medial raphe nucleus is necessary for task engagement (Ahmadlou et al., 2025). Baseline pupil size, a non-invasive readout of global arousal (Cazettes et al., 2021; Joshi & Gold, 2020), is also predictive of state transitions. Baseline pupil size is increased prior to a switch from exploitation to exploration in a bandit task in humans (Jepma & Nieuwenhuis, 2011) and mice (Shourkeshti et al., 2023). Across-trial variability of pupil size and other arousal measures (face motion energy, locomotion speed) in mice also decreases as they transition from the disengaged to engaged state, and increases when transitioning to the engaged state (Hulsey et al., 2024).

This arousal-related neuromodulatory activity may thus be an important driver of neural systems transitioning into new dynamical regimes. It has been proposed that activity of neuromodulatory regions can trigger largescale reorganisations of cortical networks (Bouret & Sara, 2005; Sara & Bouret, 2012), allowing the brain to transition to states that were previously inaccessible (Shine, 2023). To understand these dynamics, we follow a growing tradition in computational neuroscience of modelling neural systems as an attractor landscape. Behavioural states, like engagement and disengagement, can be conceptualised as “wells” that pull neural trajectories towards them over time (Guarino et al., 2025; Richman et al., 2023; Taylor et al., 2024).

Here, we investigate the relationship between arousal and engagement state during perceptual decision-making using a dynamical systems model. We build on a literature that has characterised behaviour in a rodent model of decision-making in detail, taking advantage of the large open-access dataset provided by the International Brain Laboratory (The International Brain Laboratory et al., 2021, 2025). We applied hidden Markov models to whole session time courses of response times (Ashwood et al., 2022; Gunawan et al., 2022; Hulsey et al., 2024; Kucharský et al., 2021; Kunkel et al., 2021; Visser et al., 2009), to obtain a trial-by-trial estimate of animals’ behavioural state, which we interpret as reflecting periods of engagement and disengagement in the task. We provided a conceptual replication of previous findings of changes in baseline pupil variability during state transitions (Hulsey et al., 2024), and found that baseline pupil size increases prior to transitions into the disengaged state. Using a dynamical systems framework, in which arousal acts to shape the cortical dynamics governing behavioural state, we clarify the computational role of neuromodulatory activity in behavioural state transitions.

## Results

### Identifying engagement states with hidden Markov models

95 mice (194 sessions) performed a visual decision-making task (The International Brain Laboratory et al., 2021), in which a Gabor was presented on the left or the right of the screen (Figure 1A). We define response time (RT) as time elapsed between the presentation of the stimulus and the response being registered (i.e. left or right) - note that this is after the time of first movement, which in this task is sometimes called reaction time (The International Brain Laboratory et al., 2025; Yin & Hiratani, 2025).

**Figure 1.**
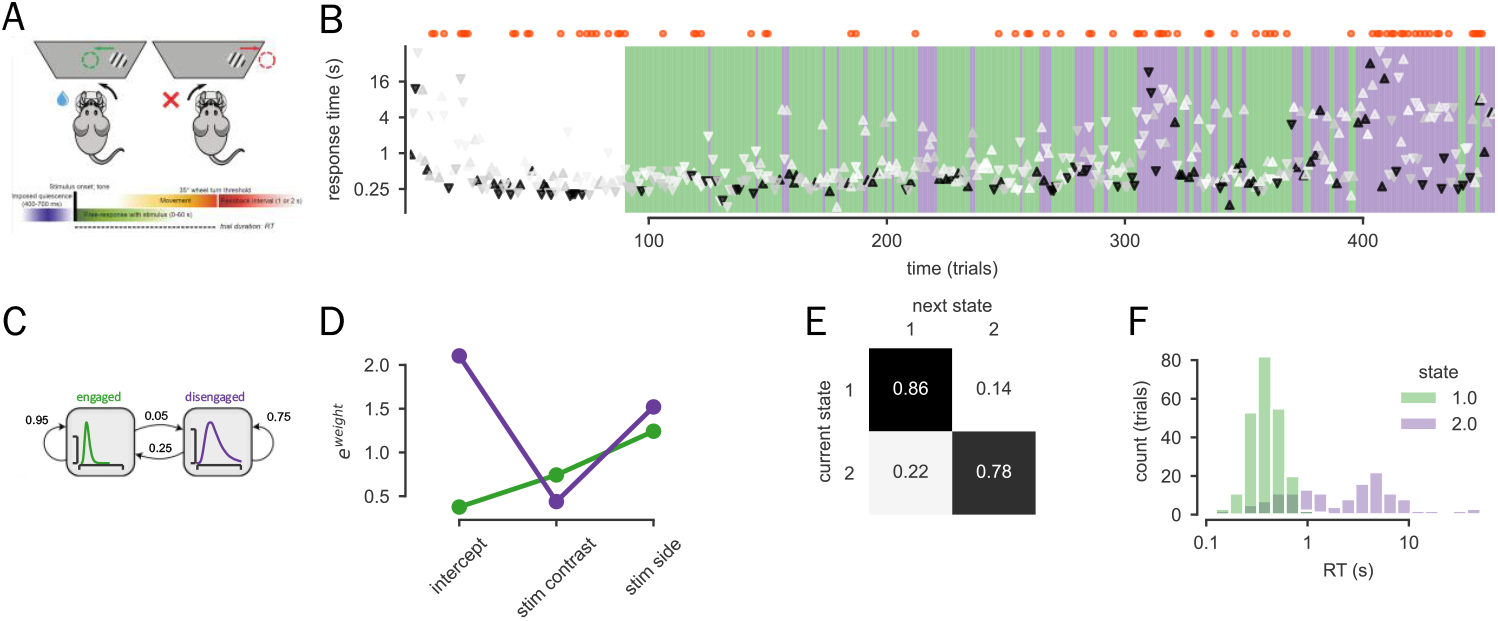
Hidden Markov Models uncover states in mouse decision-making behaviour. (A) Task design. Mice respond to a grating presented on the left or right of the screen, presented at a range of contrast levels. Response time (RT) is defined as the time from stimulus onset to response completion. Adapted from (The International Brain Laboratory et al., 2021). (B) Whole session time-course for one example session. In the foreground are the choices of the animal (left - down / right - up), with RT on the y-axis. The contrast of the presented stimulus is shown by the opacity of the arrows. The background colours show the most likely state for each trial (short-RT - green / long-RT - purple). Above the graph, red dots mark incorrect responses. The HMM was fit from trial 90 onwards (see Methods). (C) Schematic of Hidden Markov model of response times. (D) Estimated regression parameters for each state in the example session. (E) Estimated transition matrix of the example session. (F) RT distributions conditioned on state for the example session.

In order to decompose trial-by-trial behaviour into latent engagement states, we fit a two-state hidden Markov model to whole session time courses of log-transformed response times (Figure 1C). We first show model fits for one example session. Figure 1B shows the whole session of choices and response times, with the computed most likely state indicated by the background colour. We identified a fast state (green) and a slow state (purple). In this session, the mouse spent more time in the fast state. Each state was defined by a linear regression predicting log(RT) from stimulus contrast and stimulus side. We expected increases in stimulus contrast to lead to decreases in RT, while stimulus side was included because some mice show faster responses for one side than the other, possibly due to differences in visual acuity or motor preferences. The state-specific parameters of these two regressions can be seen in Figure 1D. On the y-axis is e^weight^: the effect of each variable can be interpreted as a multiplicative effect on RT (<1 indicates faster RT, >1 indicates slower RT). The intercept reflects the baseline RT with 0 contrast and a stimulus on the left. The slow state was characterised by a slower baseline RT and a greater effect of stimulus contrast and stimulus side (Figure 1D). The fast state was also slightly more ‘sticky’ than the slow state, as the probability of self-transitions was higher (Figure 1E). Finally, Figure 1F shows the distribution of RTs in each state. The slow state captures the tail of the RT distribution, with a higher mean and variance than the fast state.

HMM states were consistent across all sessions. We fit this model independently on all sessions and found that the BIC of the 2-state model was always lower than a 1-state model, supporting the existence of two generative processes underlying the session time courses of RTs (Supplementary Figure 1). The overall pattern of regression parameters mirrored that in the example session shown above: we observe one state defined by a higher intercept and stronger negative effect of contrast on RT (contrast coefficient <1; Figure 2A). This slow state captures the long tail of the RT distribution (Figure 2B), also visible in group-averaged, state-conditioned chronometric functions (Figure 2C).

**Figure 2.**
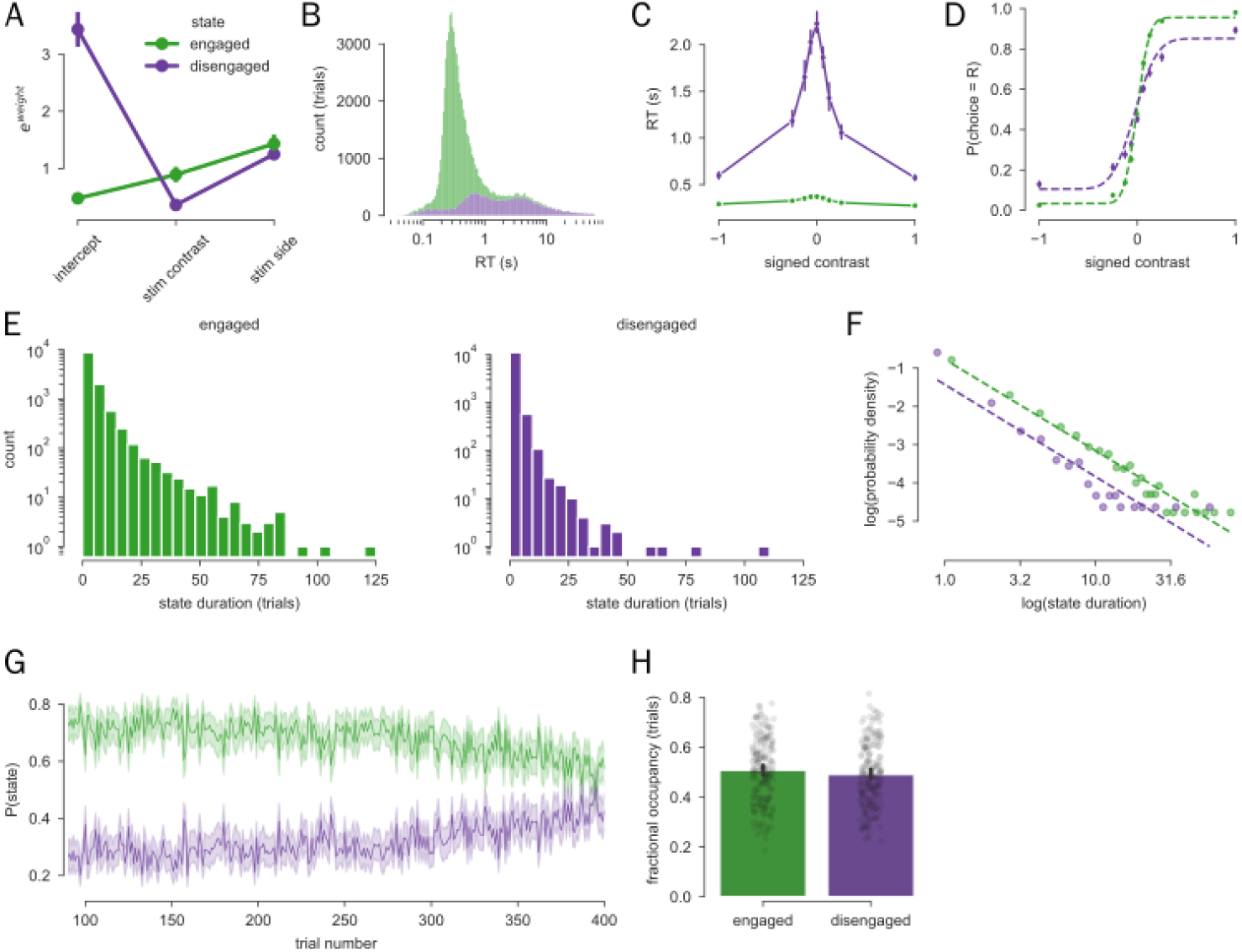
HMM fits on all 195 sessions. (A) Group-average estimated regression parameters. (B) State-conditioned RT distributions. (C) Group-averaged state-conditioned chronometric functions, showing median RT. (D) Group-averaged state-conditioned psychometric functions. Data is shown as points with error bars; dashed line is the psychometric curve fit on the average data. Statistics was conducted at the session-level. (E) Distribution of state durations, measured in trials (left - engaged, right - disengaged). (F) Power-law plots of state duration. Dashed line shows a fitted regression line (engaged - STATS; disengaged - STATS) (G) Probability of being in each state over 400 trials. (H) Fractional occupancy of each state.

To validate that the identified states capture the latent *engagement* of the animals, beyond just fast and slow RTs, we investigated the link between inferred state and accuracy, an independent measure of behavioural performance. Figure 2D shows state-conditioned psychometric curves. For each session, we fit a psychometric curve with bias, threshold and low and high lapse parameters (The International Brain Laboratory et al., 2021), and tested whether these parameters differed between states. There was a significant difference between the fast and slow states in the threshold (coef_state_=-10.99, se=0.932, z=-11.796, p=4.08×10^−32^), both lapse parameters (low: coef_state_=-0.054, se=0.007, z=-7.53, p=5.08×10^−14^; high: coef=- 0.052, se=0.007, z=-7.77, p=7.81×10^−15^), and overall accuracy (controlling for stimulus contrast, z(94)=12.86, p=1.97×10^−22^), but not in the bias parameter (coef_state_=0.58, se=1.28, z=0.477, p=0.64). This led us to conclude that the fast (green) state reflected task engagement, when animals responded quickly and accurately, while the slow (purple) state reflected disengagement, when animals responded more slowly and less accurately. This interpretation is supported by the increase in the probability of being in the slow state over the course of a session, as animals become sated and/or fatigued (Figure 2G; coef_trial_=-0.0005, se=6.75×10^−5^, z=-6.87, p=6.64×10^−12^).

Animals tended to dwell longer in the engaged state than in the disengaged state (FIgure 2E, Supplementary Figure 2). Importantly, state duration distributions appear to follow a power-law (‘1/f’) distribution, rather than the geometric distribution that would be expected from HMM transition matrices (Figure 2F). This is suggestive of specific underlying mechanisms that we later implement in a dynamical systems model. Animals spent similar proportions of the session in each state (Figure 2H; occupancy_state1_=0.5: coef=0.0092, se=0.011, z=0.825, p=0.41).

### Intermediate pupil-linked arousal is associated with highest probability of engagement

To investigate the relationship between arousal and engagement state, we used pupil diameter as a proxy of global arousal level. Pupil dilates in response to stimulus onset (Figure 3B) and feedback (Supplementary Figure 3), but also fluctuates over the course of a session, as can be observed in the ten trials shown in Figure 3A. In order to extract an estimate of baseline pupil size without contamination from the previous trial, we regressed the stimulus contrast and feedback of the previous trial out of the baseline measurement (mean pupil size from −0.25 to 0s before stimulus onset).

**Figure 3.**
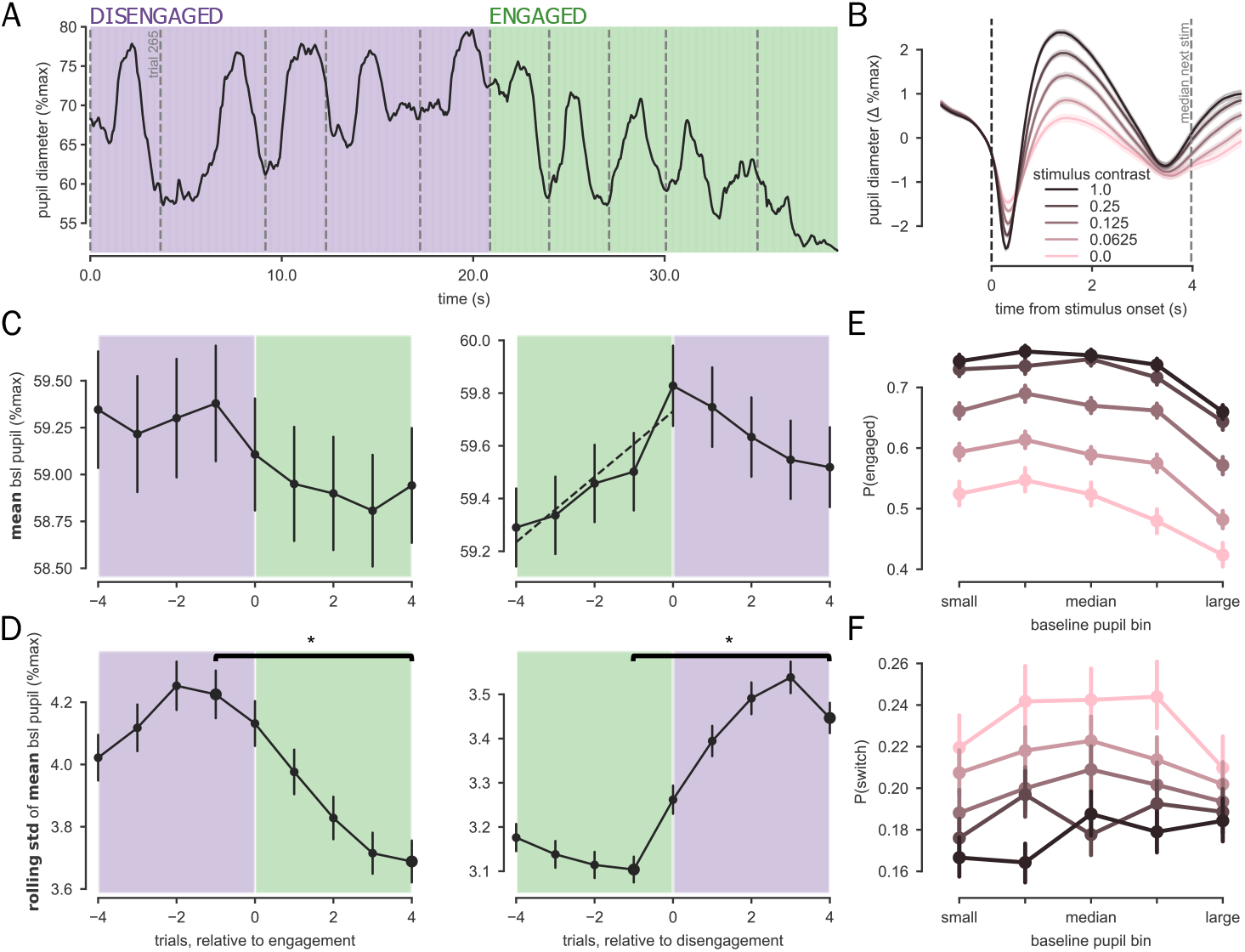
Changes in baseline pupil precede state transitions. (A) Snippet of pupil data from ten trials of the example session. Vertical, dashed, grey lines show time of stimulus onsets. The background colour reflects the inferred state at each timepoint, assuming that state transitions occur at stimulus onset. (B) Pupil dilation evoked by the presentation of the stimulus at each contrast level. Pupil time courses have been baseline corrected using a baseline period of -.25 and 0s. Pupil diameter is expressed as the change in percentage of maximum pupil diameter with respect to this baseline period. The vertical, dashed, black line marks stimulus onset, while the grey line marks the median onset time of the following stimulus (across all sessions). (C) Mean baseline pupil size surrounding a transition from the disengaged to engaged (left) and the engaged to disengaged (right) states. Background colour marks the inferred state of each trial. The dotted line in the right panel marks the significant increase in pupil size preceding a transition into the disengaged state. Only transitions following states with a duration of four trials or more are included in plots and statistics. (D) As C, with y-axis showing the standard deviation of mean pupil size over the previous four trials. Statistical tests compared the last four trials of the previous state with the first four trials of the next state; relevant points are marked by larger markers. (E-F) Probability of being in the engaged state (E) and probability of switching between states (F) as a function of baseline pupil size (x-axis) and stimulus contrast (pink - low contrast > black - high contrast).

Highest probability of being engaged was associated with intermediate baseline pupil size (Figure 3E). There were significant linear and quadratic effects of baseline pupil diameter on the probability of being in the engaged state and baseline pupil diameter, controlling for stimulus contrast (linear: t(94)=4.18, p=6.47×10^−5^; quadratic: t(94)=-4.93, p=3.48×10^−6^). This replicates many studies demonstrating a so-called inverted-U relationship between task performance and baseline pupil size (Beerendonk et al., 2023; de Gee et al., 2024; Hulsey et al., 2024; Liu et al., 2021; McGinley et al., 2015; Schriver et al., 2018; Unsworth & Robison, 2016; van den Brink et al., 2016) and suggests a relationship between (pupil-linked) arousal and behavioural state dynamics. Plots showing the relationship between baseline pupil size and RT/accuracy can be found in Supplementary Figure 3. We further investigated whether there was a relationship between baseline pupil diameter and the probability of switching between states (Figure 3F), and found no significant relationship (linear: t(94)=-1.00, p=0.32; quadratic: t(94)=1.00, p=0.32), indicating there is not a specific level of arousal at which state transitions occur.

Next, we investigated the temporal relationship between state transitions and fluctuations in arousal. While there was no consistent pattern of baseline pupil preceding switches into the engaged state (coef_trial_=-0.030, se=0.059, z=-0.508, p=0.612), there was a significant, linear increase in baseline pupil size in the four trials preceding a switch into the disengaged state (coef_trial_=0.12, se=0.027, z=4.59, p=5.38×10^−6^; Figure 3C). This contrasts with abrupt changes in stimulus contrast and accuracy preceding state transitions (Supplementary Figure 4): the linear increase remained significant when contrast and feedback preceding transitions were included in the model (coef_trial_=0.14, se=0.029, z=4.82, p=1.44×10^−6^). Across-trial variability of pupil size changes as animals transition between states. Across-trial standard deviation of baseline pupil size was higher in the last four trials of disengaged states than the first four trials of engaged states (coef_trial_=-0.53, se=0.088, z=- 6.08, p=1.15×10^−9^), and lower in the last four trials of engaged states than the first four trials of disengaged states (coef_trial_=0.34, se=0.049, z=6.99, p=2.67×10^−12^; Figure 3D) replicating previous results (Hulsey et al., 2024). This suggests that some information in baseline pupil is predictive of upcoming state transitions, implicating fluctuations in arousal as a possible driver of behavioural state transitions.

### A mechanistic model of engagement and arousal

We next constructed an attractor landscape model to understand how engagement and arousal interact and give rise to the patterns we observe in the data (Figure 4). The activity of a population of neurons can be modelled as existing in a low-dimensional state-space, which tends towards certain locations or ‘attractors’ over time. Specifically, in our model, the shape of the landscape is controlled by two parameters, α and β.The shape of the energy landscape is given by equation 1, with α determining the relative depths of the two wells, and β determining the amount of energy required to cross from one state to the other. The movement of the population activity through these states can be visualised as a ball rolling around on the landscape, the dynamics of which is given by equation 2. The activity of the system at any given time is determined by the position of this ball – at every timepoint the ball will roll downhill (the system is pulled towards a low energy state - drift), but also be injected with some random noise (diffusion), due to surrounding neural activity. The relative contribution of the noise compared to the gradient is given by *σ*. This noise can be conceptualised as faster-timescale activity of nearby neurons that interact with the population of interest (Richman et al., 2023).

**Figure 4.**
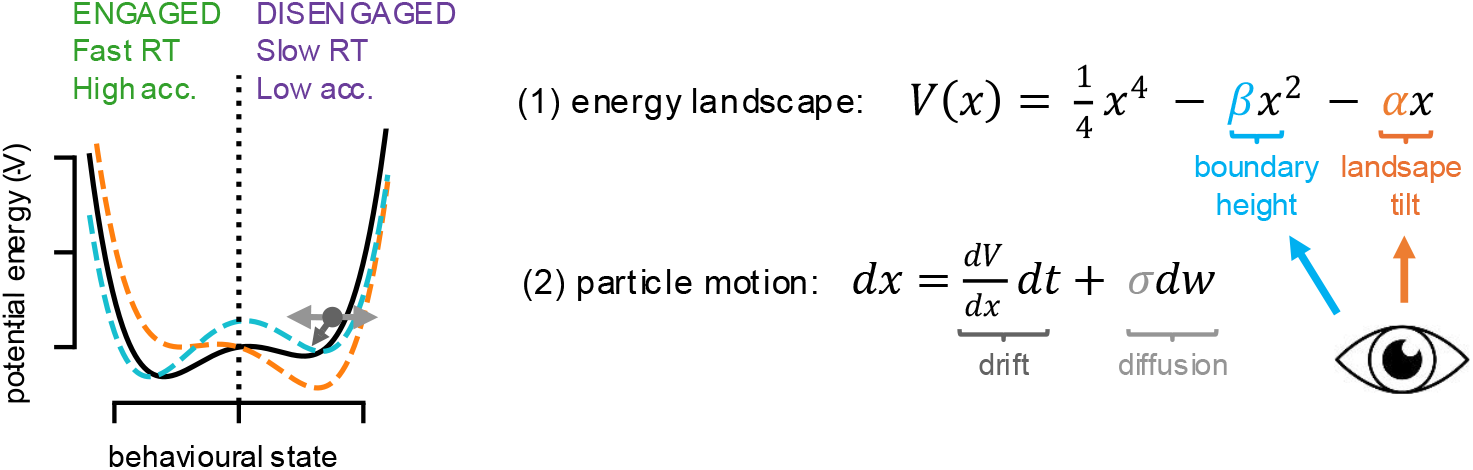
Attractor landscape model of engagement. In this model, we assume that there is a population of neurons that determine the behavioural state of an animal. The activity of this population can be modelled as an attractor landscape, with two stable steady states (“attractors” or “wells”) corresponding to engaged (left) or disengaged (right) behaviour.The orange dashed line on the landscape shows the effect of increasing α on the shape of the landscape (tilt towards the other well), while the blue dashed line shows the effect of increasing β (higher barrier between wells). We propose that arousal influences α and/or β (i.e. functioning as a control variable of the system), thereby affecting the probability of being in each state and transitioning between them.

If this model has static parameters, it generates state duration distributions which follow an exponential distribution, rather than the power-law distribution observed in our data (Figure 2F). However, introducing a stochastically fluctuating boundary height is enough to obtain power-law distributions (Mejias et al., 2010). Here, we propose that arousal, which slowly fluctuates over trials, may drive the two parameters that govern the shape of the attractor landscape that produces engagement state dynamics. This would explain why we observe changes in baseline pupil size preceding and during state transitions (Figure 3C, D), as well as the observed 1/f distributions of state duration (Figure 2F). Overall, this model provides possible explanations for observed empirical effects and a framework for understanding the relationship between engagement and arousal.

## Discussion

Most analysis of decision-making treats choices as produced from a fixed process or strategy, ignoring the temporal structure underlying sequences of decisions (Duffy et al., 2025; Urai, 2024). Studying across-trial choice variability can provide insight into the dynamics of underlying neurophysiological systems. Here, we decompose behaviour into latent engagement states. We show that mice transition between two states within sessions of decision-making, which correspond to engaged (fast and accurate) and disengaged (slow and inaccurate) behaviour (Ashwood et al., 2022). We replicate previous findings of changes in variability of baseline pupil as animals transition between states (Hulsey et al., 2024), with different data, model and an increased sample size. We also find an increase in mean baseline pupil prior to transitions into the disengaged state. This mirrors previous findings of pupil changes prior to a switch into an exploratory state during a bandit task (Jepma & Nieuwenhuis, 2011; Shourkeshti et al., 2023). While explore/exploit states do not necessarily directly map to engaged/disengaged states, task disengagement can be seen as exploring for alternative internal or external rewards (Ebitz et al., 2019; Jepma et al., 2018). We provide a conceptual model of how fluctuations in arousal could cause transitions in engagement state. Fitting this model to data, namely state duration distributions, will next enable identification of the key parameters that mediate the effect of arousal on engagement, and lead to novel predictions that can be empirically tested.

While we here take pupil size as a non-invasive proxy of the global neuromodulatory activity, direct neural recordings would enable more precise tracking of specific systems. Previous research has demonstrated that activity of acetylcholine and noradrenaline can shape cortical and subcortical dynamics (Grimm et al., 2024; Munn et al., 2021; Shine, 2023; Taylor et al., 2024) making these neurotransmitters likely candidates for influencing neural systems and behavioural state.

In conclusion, we have shown that switches between behavioural states, reflecting task engagement and disengagement, can be captured by a hidden Markov model of response times. State transitions are related to fluctuations in arousal, suggesting that arousal drives changes to the neural systems governing behaviour.

## Methods

### Sample

Data is from the International Brain Laboratory (IBL), a large public dataset of behavioural and neural data from mice performing a perceptual decision-making task. Sessions were selected from the brain-wide map data release (The International Brain Laboratory et al., 2025). Mice were fully trained on the task, and electrophysiological recordings were being completed during sessions. From this sample, sessions were excluded if any response times were negative (i.e., the mouse responded before the stimulus was presented, n = 5), pupil data could not be downloaded (n=15), or there was no significant modulation of pupil size by stimulus contrast (n=23; see Pupil Preprocessing). The resulting sample included 95 mice, 194 sessions.

### Task

The IBL task is described in detail in previous work (The International Brain Laboratory et al., 2021). In short, a Gabor was presented on the left or right of the screen, and mice responded by turning a wheel to bring the stimulus into the centre of the screen. If the wheel was moved in the correct direction (i.e. clockwise if the stimulus was presented on the left), a water reward was delivered; otherwise, a white noise pulse was presented for 500ms, followed by a 2s timeout. The next trial began after a variable quiescence period (400-700ms, randomly drawn from an exponential distribution with a mean of 550ms). The stimulus contrast was sampled uniformly from five contrast levels (100%, 25%, 12.5%, 6.25% and 0%). In the first 90 trials of all sessions, the stimulus had 0.5 probability of appearing on the left or right. For the rest of the session, the stimulus switched uncued between 0.2 and 0.8 probability of appearing on the left. These ‘bias blocks’ lasted for 0 to 100 trials (drawn from a truncated geometric distribution, on average 51 trials per block). For all analyses based on the hidden Markov model, the first 90 trials and any trials beyond 590 were excluded. The block structure is introduced after 90 trials, so starting the model fits after this avoids needing to account for a change in the structure of the task, as well as removing any ‘warm-up’ time needed by the animals. Trials beyond 590 were excluded to roughly standardise the number of trials per session. The mean number of trials analysed per session was 449.72 (se=3.18, min=311, max=500 trials).

As the response is a wheel movement rather than a button press, it is not immediately clear which time point should be taken as the moment a response is given. There are multiple timepoints of interest that correspond to different aspects of the decision-making process. Here, we decided to use the time point at which the response is registered, which is when the wheel has been moved enough to bring the stimulus across the midline of the screen (35°). We used this, as opposed to the time of the first movement of the wheel, as the duration of wheel movements varied across trials. The time of crossing the response threshold, hereafter the response time, takes into account any changes of mind or hesitations while the response is being executed (note that other studies using IBL data instead consider the time of movement initiation (Ghani, 2025; Yin & Hiratani, 2025)).

### Behavioural analysis

#### Psychometric curves

Psychometric curves (Figure 2b) were fit using the *brainbox* function *compute_psychometric*, which uses *psychofit*.*mle_fit_psycho* to fit a function with a high and low lapse, bias and threshold parameters (The International Brain Laboratory et al., 2021).

#### Hidden Markov Model

We used a hidden Markov model (HMM) to identify engagement states. The response time at trial *t* depends on a latent state *z*_*t*_ ∈ *Z*. The state space *Z* is assumed to be composed of *n* distinct states. For example, in a model with two states, this set contains the states [1, 2]. The latent state on a given trial *z*_*t*_ depends only on the state at the previous trial

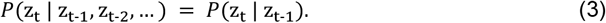

The probability of transitioning between states is described by a transition probability matrix, *A* of size *n* by *n*, where *A*_*ij*_ is the probability of transitioning from state *i* to state *j* on a given trial.

In a basic Gaussian-HMM, each latent state is associated with a Normal distribution of observations, defined by a state-dependent mean and standard deviation. However, in our case, observations were whole session time courses of response times, which depended not only on the latent state of the animal, but also on other covariates: stimulus contrast and the side of the screen on which the stimulus was presented. Stimulus contrast affects the difficulty of the task: response times were expected to be longer when contrast was lower. In some sessions, mice exhibited faster response times for one direction than the other. This side bias likely emerged as a result of motor, visual acuity or attentional biases.

In order to better fit a Normal distribution, response times were log-transformed before fitting the model: observations *s*_*t*_ *ln RT*_*t*_. We therefore defined each state using a linear regression model with *s*_*t*_ as the dependent variable and two independent variables, stimulus contrast and stimulus side:

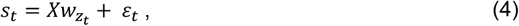

where *X* is a design matrix containing an intercept, stimulus contrast and stimulus side, and *ε*_*t*_ ∼ *N*(0,1). Note that each parameter depends on the state, *z*, so, in a two-state model, six regression parameters (three for each state), will be estimated. An HMM with linear regression observations is also referred to as a switching linear regression model.

Free parameters of the model were: the initial distribution of states at trial *t*_0_, the transition matrix *A*, and the weights of the state-dependent regression models, *w*_*z*_. HHMs were fitted in Python using the IOHMM package (https://github.com/Mogeng/IOHMM/), which applies the Expectation-Maximisation algorithm (Baum et al., 1970). This algorithm iteratively finds the maximum likelihood parameter estimates of a model, described in more detail here (Ashwood et al., 2022). In short, the algorithm estimates the most likely latent states for each trial based on the current parameter estimates (E-step), then maximises the model parameters given the expected state assignments (M-step). This process continues iteratively until convergence. This fitting algorithm guarantees that a local minimum is found, but not a global minimum. To address this, each model fit was initialised 20 times with random starting parameters. Once the model has been fit, it is possible to use the fitted parameters to estimate the posterior state probabilities of a mouse being in each state on a given trial. We assigned trials to each state by taking the most likely state.

In HMMs, the number of states is decided prior to model fitting. Different numbers of states offer different levels of detail of the dynamics underlying behaviour (Vidaurre, 2018). We decided to use a model with two latent states, as we were interested in differentiating between engaged and disengaged trials; the psychological interpretation of a three state model is unclear. Additionally, previous research applying a generalised linear model - hidden Markov model (GLM-HMM) showed that a three state model best described behaviour (Ashwood et al., 2022), where one state was interpreted as ‘engaged’ and the other two can be interpreted as ‘disengaged’ (characterised by poor sensitivity or biased responses). All disengaged states, regardless of the particular pattern of responding, were associated with longer response times than the engaged state, and can therefore be grouped as one disengaged state in our response time-based model. To ensure states were comparable across sessions and animals, we assigned state 1 as the state with the smallest intercept.

#### No response trials

If no response was given within 60s after stimulus onset, the trial timed out. The model could not be fit with NaN inputs, so we replaced any no response trials with the maximum recorded response time from the session. We also confirmed that the model fitting was robust to a different choice, namely replacing NaNs with 60s, and found that estimated parameters and state assignments were almost exactly the same.

### Pupil analysis

During experimental sessions, cameras recorded the left of the animal’s face at 60 z. Pupil diameter was estimated using the Lightning Pose algorithm (Biderman et al., 2024). Pupil diameter was downsampled to 25 Hz. Missing data up to 1s was linearly interpolated. Horizontal and vertical eye movements were linearly regressed out of the continuous pupil time course, to account for the pupil foreshortening error (Hayes & Petrov, 2016). Pupil size was expressed as a percentage of the maximum recorded pupil size of that session. To ensure high quality pupil measurement, we selected sessions in which the mean evoked pupil response from 1-2s after stimulus onset was significantly modulated by stimulus contrast.

Baseline pupil is defined as the mean pupil size from 250ms to 0ms prior to stimulus onset. There were 2.60s to 65.56s (median 4.09s) between two stimulus presentations, so often not enough time for pupil size to return to baseline after the stimulus or feedback evoked response (see Figure 3B and Supplementary Figure 3). We therefore linearly regressed out the contrast and feedback of the previous trial from the baseline pupil estimates.

## Statistical analysis

Unless stated otherwise, statistical significance was tested using generalised estimating equations (GEEs: *statsmodels*.*generalized_estimating_equations)*. This approach allows estimation of population effects, while taking into account the nested structure of data, by assuming correlation between datapoints from the same mouse. The correlation structure within mice was assumed to be exchangeable (the same across mice). We report coefficients, their standard error, z-values and p-values. Because Binomial distributions are not implemented in GEEs in *statsmodels*, statistical models of categorical data (e.g., state, feedback) or probabilities (e.g., probability of being in state 1) were estimated at the subject level, then a t-test was conducted to test whether parameters were different from zero at the group level. We report t-statistics with degrees of freedom and p-values. The threshold for significance *α* was set at 0.05. Error bars in all figures show standard error of the depicted estimator.

## Data and code availability

Data has been made available by the International Brain Lab (https://www.internationalbrainlab.com/data). Code will be shared on GitHub.

## Acknowledgements

This work was supported by the Netherlands Organisation for Scientific Research (Veni fellowship VI.Veni.212.184 to AEU, SSH-XS 406.XS.24.02.032 grant to AEU and PAJ, Vici grant VI.C.181.032 to SN).

## Supplementary Figures

**Supplementary Figure 1.**
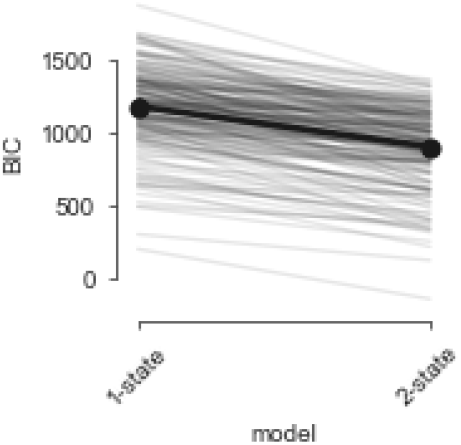
Bayesian Information Criteria calculated for a 1-state and 2-state logistic regression model. Individual sessions are shown in grey; group average is in black.

**Supplementary Figure 2.**
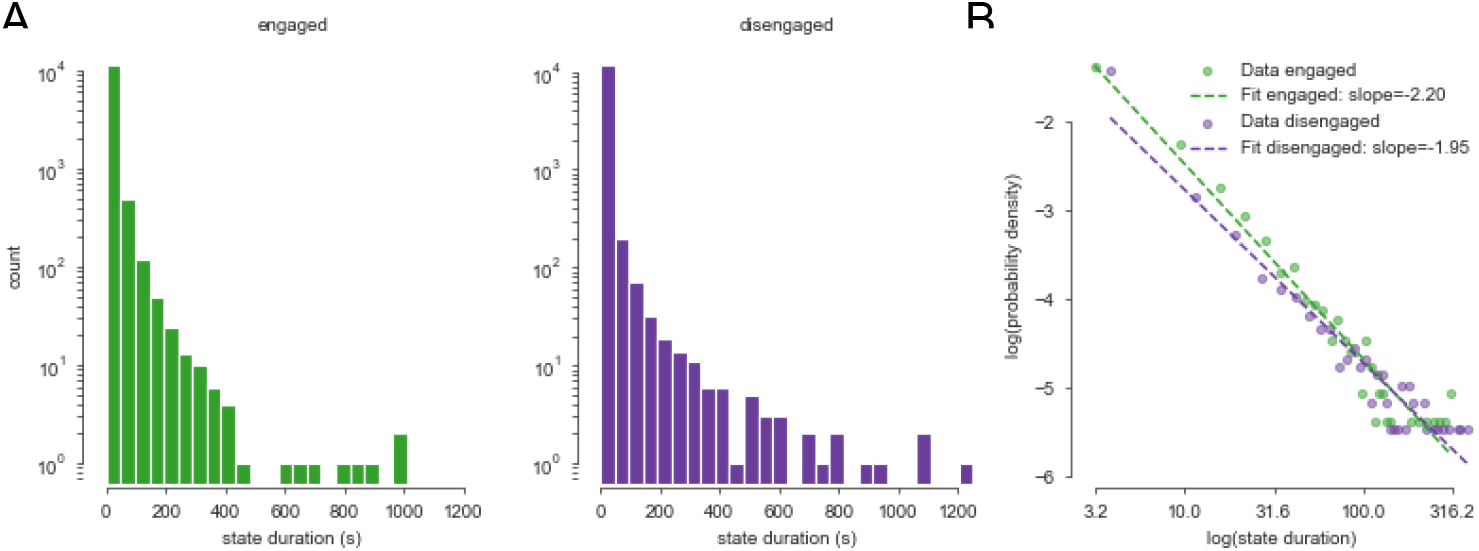
Dwell times (A) and power law plot (B) with state duration measured in seconds. **Calculating the state durations in seconds, by assuming that state transitions occurred at the time of stimulus onset, revealed that animals dwelled in the engaged and disengaged states for similar lengths of time - despite dwelling in the disengaged state for fewer trials in a row, RT in those trials was longer, and there were more frequent errors (Figure 2D), leading to 1s timeouts.**

**Supplementary Figure 3.**
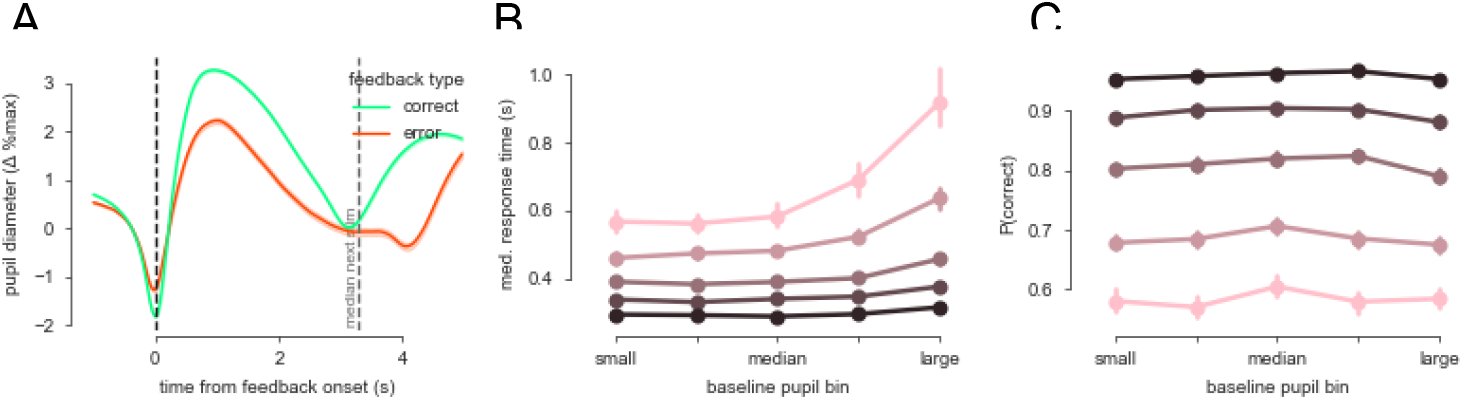
Additional pupil plots. (A) Pupil dilation evoked after correct (green) and error (red) trials. Pupil timecourses have been baseline corrected using -.25 and 0s before feedback onset. The vertical, dashed, black line marks feedback onset, while the grey line marks the median onset time of the following stimulus (across all sessions). (B-C) Median response time (B) and probability of being correct (C) as a function of baseline pupil size (x-axis) and stimulus contrast. Controlling for stimulus contrast, there was a significant quadratic relationship between baseline pupil size and response time (coef_quad_=9×10^−4^, se=4.18×10^−4^, z=2.10, p=0.36) and a significant linear and quadratic relationship between baseline pupil and accuracy (linear: t(94)=3.00, p=0.0035; quadratic: t(94)=-3.76, p=2.95×10^−4^).

**Supplementary Figure 4.**
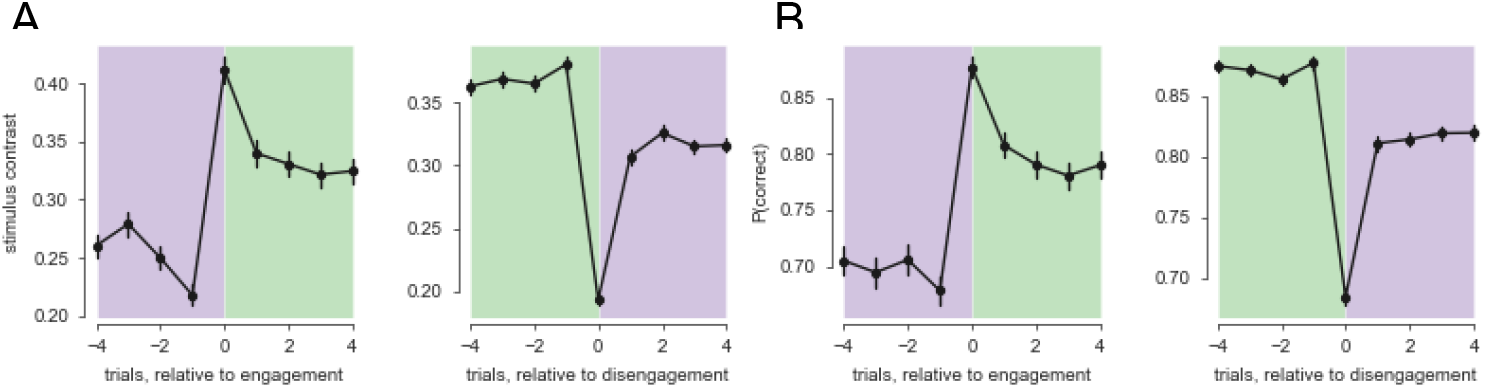
Task variable preceding state transitions. Contrast (A) and accuracy (B) in the trials surrounding a transition into the engaged (left) and disengaged (right) state. In the double-well model, we do not explicitly model the effects of task-related variables (such as stimulus contrast and reward), although it is likely that these interact with endogenous fluctuations in pupil-linked arousal. Mice are more likely to transition from disengaged to engaged after a reward, and vice versa after an error. However, we make no claim that these factors are independent. For example, if an animal has relatively high arousal at a certain timepoint, receiving a reward could ‘boost’ overall arousal and cause a transition into a disengaged state (Shourkeshti et al., 2023). In this paper, we are concerned with neural causes, rather than external causes, but both explanations can co-exist. Future work should integrate the effects of reward and other external factors into an attractor landscape model.

