## Supplementary Figures for "A dynamical systems model of arousal-driven behavioural state transitions"

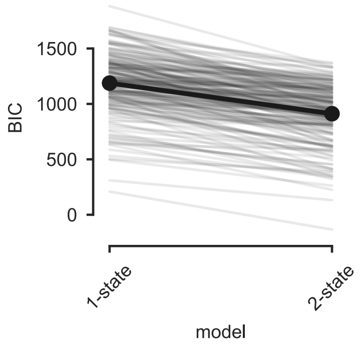


**Supplementary Figure 1. Bayesian Information Criteria calculated for a 1-state and 2-state logistic regression model.** Individual sessions are shown in grey; group average is in black.

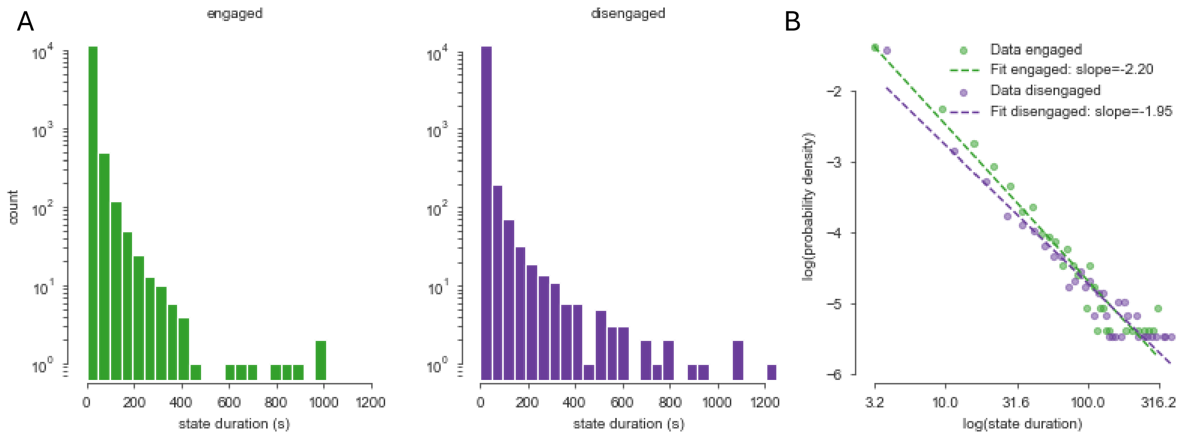


**Supplementary Figure 2. Dwell times (A) and power law plot (B) with state duration measured in seconds.** Calculating the state durations in seconds, by assuming that state transitions occurred at the time of stimulus onset, revealed that animals dwelled in the engaged and disengaged states for similar lengths of time - despite dwelling in the disengaged state for fewer trials in a row, RT in those trials was longer, and there were more frequent errors (Figure 2D), leading to 1s timeouts.

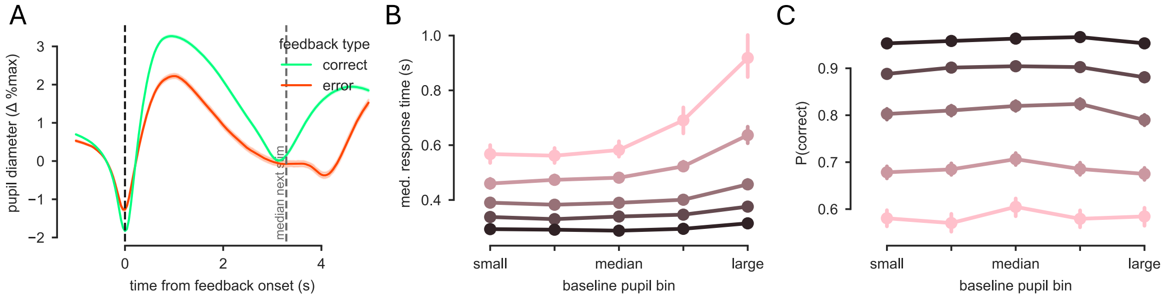


**Supplementary Figure 3. Additional pupil plots.** (A) Pupil dilation evoked after correct (green) and error (red) trials. Pupil time courses have been baseline corrected using -.25 and 0s before feedback onset. The vertical, dashed, black line marks feedback onset, while the grey line marks the median onset time of the following stimulus (across all sessions). (B-C) Median response time (B) and probability of being correct (C) as a function of baseline pupil size (x-axis) and stimulus contrast. Controlling for stimulus contrast, there was a significant quadratic relationship between baseline pupil size and response time ($coef_{quad}$=9x$10^{-4}$, se=4.18x$10^{-4}$, z=2.10, p=0.36) and a significant linear and quadratic relationship between baseline pupil and accuracy (linear: t(94)=3.00, p=0.0035; quadratic: t(94)=-3.76, p=2.95x$10^{-4}$).

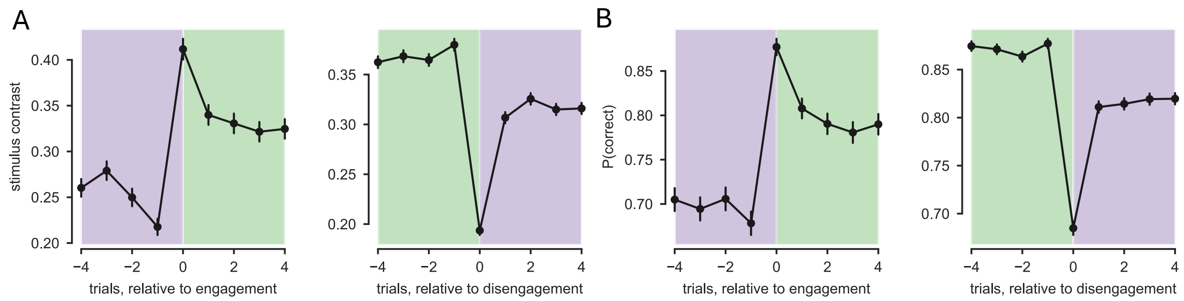


**Supplementary Figure 4. Task variable preceding state transitions**. Contrast (A) and accuracy (B) in the trials surrounding a transition into the engaged (left) and disengaged (right) state. In the double-well model, we do not explicitly model the effects of task-related variables (such as stimulus contrast and reward), although it is likely that these interact with endogenous fluctuations in pupil-linked arousal. Mice are more likely to transition from disengaged to engaged after a reward, and vice versa after an error. However, we make no claim that these factors are independent. For example, if an animal has relatively high arousal at a certain timepoint, receiving a reward could 'boost' overall arousal and cause a transition into a disengaged state (Shourkeshti et al., 2023). In this paper, we are concerned with neural causes, rather than external causes, but both explanations can co-exist. Future work should integrate the effects of reward and other external factors into an attractor landscape model.
